# Eukaryotic core histone diversification in light of the histone doublet and DNA topo II genes of Marseilleviridae

**DOI:** 10.1101/022236

**Authors:** Albert J. Erives

## Abstract

While eukaryotic and archaean genomes encode the histone fold domain, only eukaryotes encode the core histones H2A, H2B, H3, and H4. Core histones assemble into a hetero-octamer rather than the homo-tetramer of Archaea. Thus it was unexpected that core histone “doublets” were identified in the cytoplasmic replication factories of the Marseilleviridae (MV), one family of Nucleo-Cytoplasmic Large DNA Viruses (NCLDV). Here we analyze the core histone doublet genes from all known Marseilleviridae genomes and show that they encode obligate H2B-H2A and H4-H3 dimers of likely proto-eukaryotic origin. Each MV core histone moiety forms a sister clade to a eukaryotic core histone clade inclusive of canonical core histone paralogs, suggesting that MV core histone moieties diverged prior to eukaryotic neofunctionalizations associated with paired linear chromosomes and variant histone octamer assembly. We also show that all MV genomes encode a eukaryote-like DNA topoisomerase II enzyme that forms a clade that is sister to the eukaryotic clade. As DNA topo II influences histone deposition and chromatin compaction and is the second most abundant nuclear protein after histones, we suggest MV genes underlie a proto-chromatinized replisome that diverged prior to diversification of eukaryotic core histone variants. Thus, combined domain architecture and phylogenomic analyses suggest that a primitive origin for MV chromatin genes is a more parsimonious explanation than horizontal gene transfers + gene fusions + long-branch attraction constrained to each core histone clade. These results imply that core histones were utilized ancestrally in viral DNA compaction, protection from host endonucleases, and/or other unknown processes associated with NCLDV-like progenitors.

## Introduction

The Marseilleviridae (MV) are a distinct family of viruses within the Nuclear-Cytoplasmic Large DNA Viruses (NCLDV) (Yutin *et al.* 2009; Koonin and Yutin 2010; Yutin *et al.* 2014) with eukaryote-like core histone genes (Boyer *et al.* 2009; Thomas *et al.* 2011). These histone genes are unusual in at least three ways: (***i***) most of the histone domains are orthologous to eukaryotic core histones (H2A, H2B, H3), but one (h) has been weakly assigned to the single archaeal histone clade (Thomas *et al.* 2011); (***ii***) the core MV histone domains are “fused” into divergently transcribed doublet genes, thus encoding forced h-H3 and H2B-H2A heterodimers; and (***iii***) MV histone proteins were reported found in the virus particles of Marseillevirus, thus suggesting MV nucleosomes function in the compaction, protection, and/or regulation of their large viral genomes (Boyer *et al.* 2009). The MV core histone repertoire is unlike archaeal genomes (Bailey *et al.* 1999; Reeve *et al.* 2004; Ammar *et al.* 2012), which typically encode a single histone that forms homodimers and a tetramer nucleosome that protects ∼60 bp DNA (Bailey *et al.* 1999). Furthermore, archaeal histones are short peptide sequences 65–69 amino acids in length (Reeve *et al.* 2004) and lack the N-terminal histone tails of eukaryotic histones, which are heavily modified by covalent attachments of acetyl and methyl groups to specific conserved lysine residues present throughout the N-terminal histone tails, which protrude away from the histone octamer (Turner 2014). Genes for linker histones (H1/H5) have not yet been found in any of the six known Marseilleviridae genomes. These include the 368 kb Marseillevirus genome (Boyer *et al.* 2009), the 374 kb Cannes 8 virus genome (Aherfi *et al.* 2013), the 346 kb Lausannevirus genome (Thomas *et al.* 2011), the 386 kb Insectomime virus genome (Boughalmi *et al.* 2013), and the 380 kb Tunis virus genome (Aherfi *et al.* 2014a). About 300–400 genes are shared amongst the Marseilleviridae viruses, with about ∼600 protein-coding genes present in the pan-MV genome (Aherfi *et al.* 2014b).

Initial phylogenetic analysis of the Marseillevirus and Lausannevirus histones genes revealed a “challenging” perspective of MV histone origins, evolution, and relationships to eukaryotic histones (Thomas *et al.* 2011). Investigation of the origin and make-up of the MV histones might be relevant and/or informative to understanding the origin of eukaryotes, which exceed prokaryotes in cellular complexity, compartmentalization, and their sophisticated chromatinized genomes (Koonin 2010; Rochette *et al.* 2014). MV chromatin might constitute an independent model for the evolutionary chromatinization of a genome if the MV histone genes were acquired from an unknown eukaryotic group and were maintained for eukaryote-like chromatinization. Alternatively, the MV core histones might be derived from a stem-eukaryote “ghost” lineage and could provide information on the evolutionary origin of eukaryotes.

To investigate the evolutionary origins of MV histones, we consider recently sequenced MV genomes (*e.g.*, Insectomime virus/Tunisvirus, Cannes 8 virus, and Melbournevirus) discovered since the Marseillevirus and Lausannevirus reports (Aherfi *et al.* 2013; Boughalmi *et al.* 2013; Aherfi *et al.* 2014a; Aherfi *et al.* 2014b; Doutre *et al.* 2014). We also consider both canonical core histones, which are unique to eukaryotes, as well as eukaryotic variants such as H2A.Z and CenH3/CENP-A. The H2A.Z histone is associated with core promoter nucleosomes, which is likely the ancestral function across eukaryotes and Archaea (Ammar *et al.* 2012). The histone cenH3 is a fast-evolving eukaryotic histone associated with centromeric nucleosomes (Henikoff and Furuyama 2012). Centromeres are the chromosomal regions targeted by kinetochores, which are a eukaryote-specific innovation associated with their linear chromosomes and which function in proper chromosomal segregation (Henikoff and Furuyama 2012). While centromeric H3 variants are fast evolving and can be lost (Drinnenberg *et al.* 2014) and presumably replaced with co-opted H3 paralogs (Wang *et al.* 2011; Masonbrink *et al.* 2014; Yuan *et al.* 2015), inclusion of these variants could still be informative. Last, we consider eukaryotic groups among the Discoba super-group, which recent phylogenetic analysis puts as the sister group of all other eukaryotes (He *et al.* 2014). Within this group are the Excavates and in particular the kinetoplastid protists (*e.g.*, Trypanosome parasites), which lack homologous components of the kinetochore, which assembles on centromeres (Akiyoshi and Gull 2013). Several excavates have divergent H3 variants utilized in non-centromeric regions such as telomeres (Lowell and Cross 2004), polycistronic loci (Siegel *et al.* 2009), or in other non-canonical chromosomal structures (Dawson *et al.* 2007; Akiyoshi and Gull 2014).

Here we show that the core histone moieties of MV and other related genes are likely derived prior to the evolutionary diversification of eukaryotic core histone variants. First, we use phylogenetic analysis to place these core MV histone domains as sister clades to each of the four eukaryotic core histones. The fused H2B-H2A and H4-H3 configurations of these genes is understandable given that the MV H2A and H3 clades are outgroups to eukaryotic core variants (*i.e*., paralogy groups) that have evolved specialized functions in the context of the eukaryotic chromosome (intergenic DNA, centromeres, etc.). Second, we show that other MV genes encoding DNA replisome components adapted to a chromatinized template, such as DNA topoisomerase II, also form a clade that is sister to all eukaryotic DNA topo II sequences. Third, the maintenance of these core histone fusions and DNA topo II genes in divergent MV genomes suggests that these genes are under purifying selection and are not recent horizontal transmissions. Together these results suggest that further study of NCLDV genomes may be informative for understanding eukaryotic origins.

## Materials and Methods

**<u>Gene sequences</u>**. All sequences were obtained from the following genomes: Cannes 8 virus, complete genome, 374,041 bp circular DNA, Accession: KF261120.1. Melbournevirus isolate 1, complete genome, 369,360 bp circular DNA, Accession: NC_025412.1. Tunisvirus fontaine2 strain U484, complete genome, 380,011 bp circular DNA, Accession: KF483846.1. Marseillevirus marseillevirus strain T19, complete genome, 368,454 bp circular DNA, Accession: NC_013756.1. Lausannevirus, complete genome, 346,754 bp circular DNA, Accession: NC_015326.1. See text for associated publications of each genome.

**<u>Phylogenetic analyses</u>**. The MUSCLE (MUltiple Sequence Comparison by Log-Expectation) alignment algorithm and MEGA6 were used to generate alignments of the Med12 and Med15/Mdt15 sequences (Edgar 2004a; Edgar 2004b; Tamura *et al.* 2013). Phylogenetic analysis was conducted using Bayesian MCMC and mixed amino acid models were tested via MrBayes (Huelsenbeck and Ronquist 2001; Ronquist and Huelsenbeck 2003; Ronquist *et al.* 2012). Sufficient generations were run for the average standard deviation of split runs to be less than 1%. The numbers on the nodes in the Bayesian trees represent posterior probabilities. All other trees give bootstrap replicates and were computed using MEGA6 (see also figure legends).

## Results

**MV core histone moieties predate eukaryotic core histone diversification**

All Marseilleviridae genomes that have been identified contain the three histone genes originally seen in Lausannevirus and Marseillevirus: a pair of divergently transcribed histone doublets H2B–H2A and h–H3, where h is an ambiguous histone domain that groups either with the single archaeal histone clade (h), or else with the eukaryotic H4 core histone clade (Thomas *et al.* 2011). All five major MV genomes have all three genes containing domains from characteristic of the H2A-domain superfamily seen in eukaryotes and Archaea (Fig. S1). Phylogenetic analysis of a concatenated alignment between all three H2A-domain containing genes shows that they are slowly evolving (Fig. S1).

To re-evaluate the intriguing MV core histone repertoire in light of more recent MV genomes (Aherfi *et al.* 2014a; Aherfi *et al.* 2014b; Doutre *et al.* 2014), we phylogenetically analyzed the four histone domains of the obligate doublet genes (see supplementary alignment files for sequences). The H2B and h moieties of the divergently transcribed histone doublet genes each occur in the N-terminal halves of the predicted proteins (Fig. 1A). Interestingly, this leaves the H2A and H3 moieties in their respective C-terminal halves and these correspond to the core histones associated with distinct functional variants, such as H2A.Z and cenH3/CENP-A.

**Figure 1.**
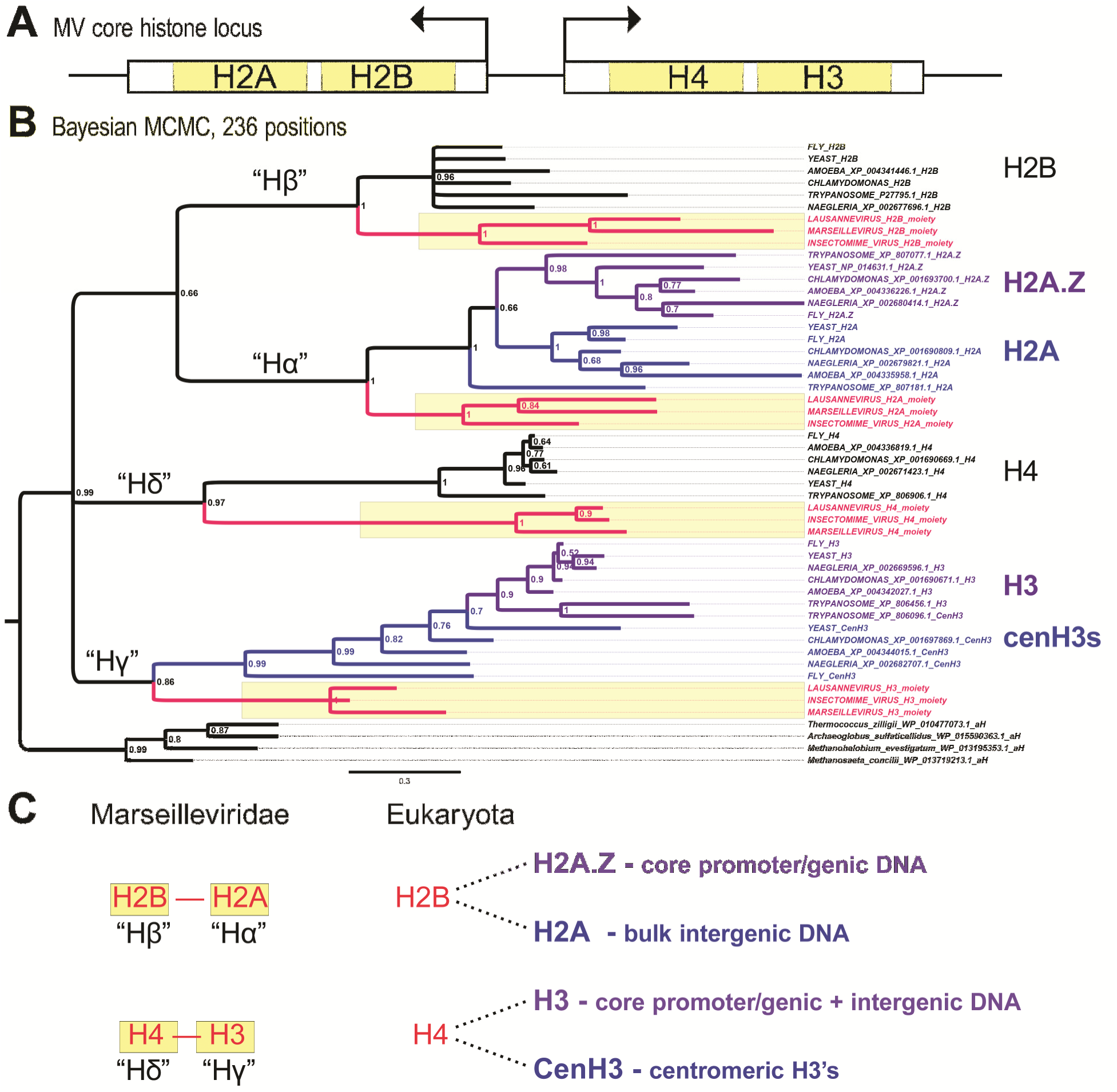
The MV core histone locus defines a full repertoire of basal eukaryotic core histones. (**A**) All MV genomes possess a pair of divergently transcribed histone doublet genes with two domains each for H2B and H2A, or H4 and H3. (**B**) Bayesian MCMC-based Inference shows that each MV core histone domain defines a well-supported sister clade (yellow highlighted clades with red lineages) to the eukaryotic core histone groups, including eukaryotic core variants for H2A.Z/H2A and H3/CenH3 (purple and blue clades within each core family). Posterior probabilities following 2,000,000 generations of sampling mixed amino acid substitution models with 25% burn-in generations and without Metropolis coupling (heated chains) are indicated at all nodes. The average standard deviation of split frequencies from two parallel runs was less than 1% (0.0076). This analysis indicated posterior probabilities of 80.5% and 19.5% for the Wag (Whelan and Goldman 2001) and Blosum (Henikoff and Henikoff 1992) amino acid models, respectively. All core histone clades from eukaryotes and MV viruses are grouped in a single super clade of the core-histones with a posterior probability of 0.99. Representative archaeal histones are grouped together in an outgroup clade at the bottom. This analysis was conducted on an alignment using MUltiple Sequence Comparison by Log-Expectation (MUSCLE) (Edgar 2004b). (**C**) The above phylogenetic analysis suggests that that the basal core histones predate eukaryote-specific duplications and neo-functionalizations in the Hα and Hγ clades. Interestingly, some of these eukaryote-specific specializations are associated with intergenic nucleosomes (eukaryotic H2As) or centromeric nucleosomes (cenH3s). Thus, the core basal histones, defined as Hα, Hβ, Hγ, and Hδ likely predate the evolutionary innovation of large, linearized chromosomes with centromeric pairing mechanisms, a late stem-eukaryotic innovation.

We conducted Bayesian phylogenetic analysis (Huelsenbeck and Ronquist 2001; Ronquist and Huelsenbeck 2003; Ronquist *et al.* 2012) based on global alignments constructed via MUSCLE, MUltiple Sequence Comparison by Log-Expectation (Edgar 2004b), and/or CLUSTALW (Thompson *et al.* 2002) of the following: (*i*) the four separated MV histone moieties (Fig. 1A) from the three most divergent MV genomes of Insectomime, Lausannevirus, and Marseillevirus, as indicated by our phylogenetic analysis of all known MV genomes (Fig. S1). We also included eukaryotic sequences for key core histone variants including H2A.Z in addition to “canonical” H2A, and cenH3 in addition to canonical H3, as well as representative histones from Euryarchaeota.

We find that the four core MV histone domains form highly-supported sister groups to the four core histone clades of eukaryotes (Fig. 1B). These results resolve that the h MV histone moiety is derived from a eukaryote-like core histone H4 super-clade and not the archaeal histone clade. Resolution of this cryptic histone domain as a *bona fide* H4 ortholog is consistent with its joined condition alongside its canonical H3 partner. Thus, the standard MV genomic configuration of a full core histone repertoire of forced H2B-H2A and H4-H3 doublets is remarkably similar to an anticipated basal/intermediate eukaryotic condition of fused histone doublets (Malik and Henikoff 2003).

There is perfect support for an ancestral eukaryotic duplication that produced H2A and H2A.Z (compare purple H2A.Z and blue H2A sister clades in Fig. 1B). As H2A.Z retains the ancestral function of participating in the +1 nucleosome of genes at their core promoters as well the downstream genic nucleosomes to a lesser extent (Ammar *et al.* 2012), it is likely that H2A is a neofunctionalized H2A.Z paralog that became specialized for bulk intergenic DNA. Given the presence of an “H2A” moiety in MV and phylogenetic principles for histone naming (Talbert and Henikoff 2013), we refer to the H2B-H2A and H4-H3 doublet genes as the Hβ-Hα and Hδ-Hγ genes, respectively (Fig. 1C). Within this framework, Hα is the ancestral core histone ortholog that duplicated in the late stem-eukaryotic lineage to give rise to distinct H2A.Z and H2A paralogs.

Histone neo-functionalization during late stem-eukaryotic evolution is also seen in the duplication of Hγ into H3 and centromeric H3s (Fig. 1B). As previously found (Malik and Henikoff 2003; Drinnenberg *et al.* 2014), support for a single clade of centromeric H3 variants is lacking (Fig. 1B). Nonetheless, the phylogenetic analysis (Fig. 1B) gives near perfect support for a well formed, and slowly-evolving, eukaryotic core-histone H3 clade, as well as a super-clade that includes the fast-evolving centromeric H3’s. This analysis provides evidence for a duplication of Hγ into a general eukaryotic H3 with an ancestral genic-marking role plus a newer intergenic role, and neofunctionalized H3 paralogs co-opted into centromeric roles (Fig. 1C). These phylogenetic results suggest that the MV core histones likely predate the eukaryotic innovation of linear paired chromosomes with centromeres and substantial intergenic DNA.

**MV DNA topoisomerase II is eukaryote-like but not assignable to any eukaryotic lineage.** Components of the eukaryotic replication fork also interact with conserved eukaryotic machinery functioning in histone deposition and chromatin compaction. To investigate whether additional genes are conserved across the Marseilleviridae that might function in a replisome complex adapted to working with core histone-based nucleosomes, we considered all 127 annotated Lausannevirus proteins with known domains to see which are most conserved with eukaryotes. We conducted a BLASTP query of the *Saccharomyces cerevisiae* proteome, and identified DNA topoisomerase II homolog as the best match (1e-174 E-value, 1,009 amino acid long protein encoded by yeast *TOP2*). DNA topoisomerase II is the second most abundant eukaryotic nuclear protein after histones, and influences chromatin compaction and histone deposition (Roca 2009; Nikolaou *et al.* 2013). The major domain of eukaryotic DNA topoisomerase II is homologous to the archaeal DNA gyrase subunit B, which can provide an outgroup sequence.

We conducted a Maximum Likelihood analysis of all alignment columns with ≥ 90% data (Fig. 2A), as well as Bayesian MCMC analysis (Fig. 2B). Both trees place the MV DNA topoisomerase II genes as sister group to Eukarya with Archaea as an out-group (Fig. 2). Thus, these results cannot rule out that the highly-conserved DNA topoisomerase II gene of Marseilleviridae is derived from a stem-eukaryotic lineage. To further test the idea that the MV topoisomerase II gene is not derived from a specific extant eukaryotic lineage, we used the predicted Lausannevirus Topo II protein sequence to search for the most similar eukaryotic homologs. The top eleven hits are fungi and have nearly perfect query coverage over the nearly 1,200 residues with about 37%–35% amino acid identity. We then subjected a new alignment with only the eukaryotic and MV taxa to Bayesian phylogenetic analysis. This analysis shows that despite the closest similarity to some fungal DNA topo II peptide sequences, the MV topoisomerase II genes were not derived from any one eukaryotic clade (Fig. S2).

**Figure 2.**
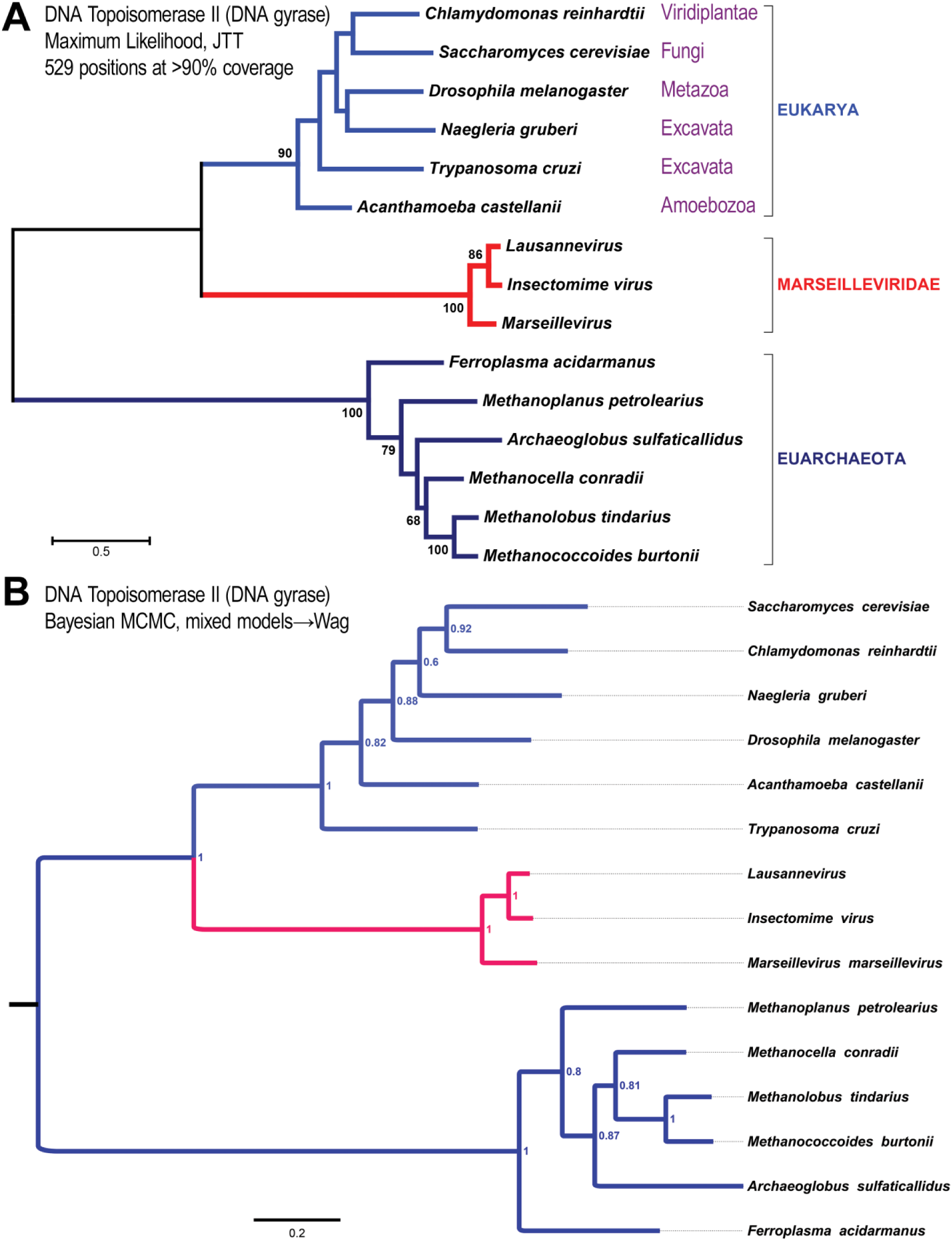
All MV genomes encode a basal eukaryotic DNA topoisomerase II protein. The presence of eukaryote-like nucleosomal chromosomes in MV is likely to require adaptations in the complexes operating at viral DNA replication forks. Analysis of MV genomes shows that the most conserved gene across MV and Eukarya is the gene encoding the large DNA topoisomerase II enzyme, which functions at the replisome. Together, phylogenetic analyses of the MV core histones and the MV DNA topoisomerase II show that these MV genes do not group within eukaryotes or Archaea. (**A**) Phylogenetic analysis by Maximum Likelihood on an alignment including all 529 columns with >90% data from all taxa and the JTT substitution matrix(Jones *et al.* 1992). Support values are from 500 bootstrap replicate data sets from 529 alignment columns that remained after a threshold cut-off of 90% data was applied. Only percent bootstrap replicate values >60% are shown. (**B**) Phylogenetic analysis by Bayesian MCMC using the same alignment as in (**A**). 2,000,000 generations of parallel runs with final average standard deviation of split frequencies = 0.0129 with 25% burn-in generations after sampling mixed amino acid models. Final generations sampled the Wag substitution model.

## Discussion

We found phylogenetic support for the possibility that the MV core histones and DNA topoisomerase II enzyme are derived from a stem-eukaryotic lineage that predates neofunctionalization of canonical eukaryotic histone paralogs (Fig. 3, “early hypothesis”: steps 1→2→3). This stem-lineage spans several evolutionary steps leading to LECA, the Last Eukaryotic Common Ancestor (Fig. 3). The MV sequences encoding the core histone moieties and large DNA topo II enzyme appear to be derived from a point along the currently accepted stem-eukaryotic lineage. This point would bisect that lineage into an early branch defined by the evolution of the four core histones (Hα, Hβ, Hγ, and Hδ) that may have been ancestrally present as doublets, and a later lineage featuring split diversified core histones (*e.g.*, Hβ-Hα → H2B + H2A|H2A.X |H2A.Z|macroH2A, and Hδ-Hγ → H4 + H3|H3.3|cenH3). Thus, the singlet nature of these core histones likely facilitated combinatorial interactions with specialized variants for genic DNA (H2A.Z and H3), intergenic DNA (H2A), and centromeric DNA (cenH3s). Furthermore, the early branch already featured a eukaryote-like DNA topoisomerase II containing additional peptide sequence not seen in archaeal DNA gyrase.

**Figure 3.**
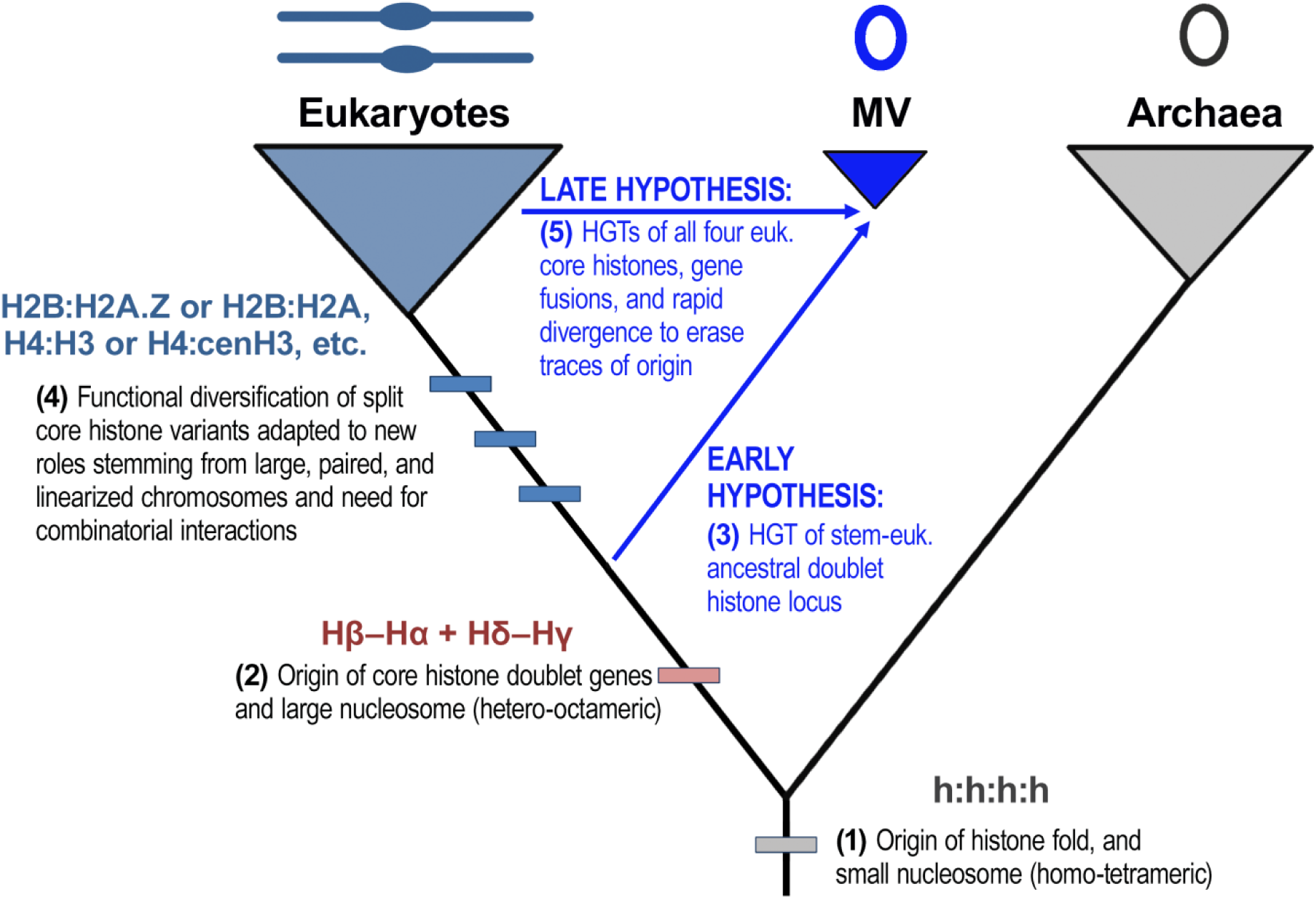
Early and late origin hypotheses for MV genes related to a chromatinized replisome. In the early hypothesis, MV core histone doublets were derived by horizontal gene transfer (HGT) from a proto-eukaryotic ancestor possessing fused core histone variants (steps 1→2→3) prior to the evolutionary diversification of eukaryotic core histone variants for H2A and H3. In the late hypothesis, MV core histone doublets were derived by HGT from eukaryotic lineage(s) after eukaryotic diversification and after the evolution of core histone variants (steps 1→2→4→5). These new HGT genes would then have fused to produce the doublet gene architecture conserved across MV.

The alternative hypothesis to an early stem-eukaryotic origin for MV genes is the “late origin hypothesis” in which an MV ancestor acquired all four core histone genes by horizontal gene transfer (HGT) from specific eukaryotic lineages, followed by subsequent core histone gene fusion (Fig. 3, steps 1→2→4→5). However, this late hypothesis would also require these genes to diverge sufficiently so as to eradicate any measurable affinities of these MV core histones to their eukaryotic source lineage. While this is quite possible for the small core histone moieties, it becomes less likely for the large and highly conserved DNA topoisomerase II protein. Thus, it would appear that the early origin hypothesis is the most parsimonious one of the two.

The configuration of a full histone core repertoire in a pair of divergently transcribed histone doublets in MV is remarkable, notwithstanding eukaryotic examples of extreme core histone divergence. One such example is the repeated loss of centromeric H3s in insect lineages with derived holocentric chromosomes (Drinnenberg *et al.* 2014). Another example is the Bdelloid rotifer class, which is the largest known clade of animals to be obligately asexual (Mark Welch and Meselson 2000; Flot *et al.* 2013). This class of rotifers substituted high molecular mass H2A variants in place of (*i*) the canonical H2A histone, which is present in nearly all eukaryotes; and (*ii*) the H2AX histone, which is involved in eukaryotic DSB repair (Van Doninck *et al.* 2009).

Recent speculation has turned to the possibility that new domains of life may be discovered in the era of massive genomic sequencing and analysis (Woyke and Rubin 2014). The absence of ribosomal RNAs disallows the ability to use such markers in the case of most viruses, although many NCLDV families possess abundant tRNA repertoires (Michely *et al.* 2013). Nonetheless, some “fourth domain” signals in specific genes of the NCLDV have been proposed (Claverie 2013; Woyke and Rubin 2014). The evidence presented here does not imply a fourth domain origin for the shared MV genome, but rather a fourth domain origin for the MV repertoire of core histone doublet and DNA topoisomerase II genes. Additional future studies might address whether the evolution of large viral genomes could have led to core histones functioning in compaction of viral DNA into capsids, and/or protection of viral DNA from prokaryotic endonucleases.

## SUPPLEMENTARY FIGURES

**Figure S1.**
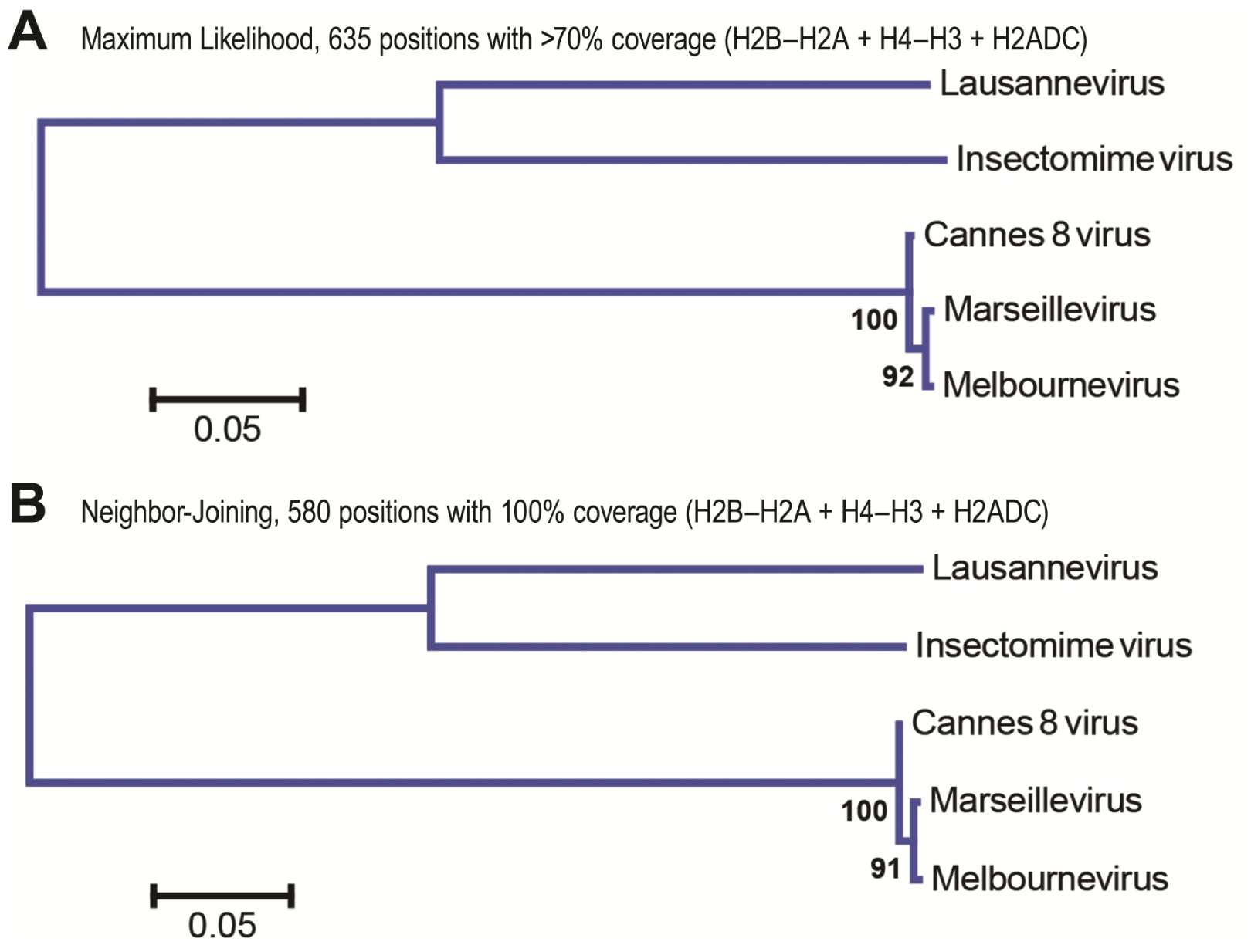
Evolution of histone genes from all known Marseilleviridae (MV). Phylogenetic analysis of all known histone-containing proteins predicted to be encoded in Marseilleviridae genomes including Lausannevirus, Insectomime virus, Cannes 8 virus, Marseillevirus, and Melbournevirus shows that these are slow evolving. (**A**). Phylogenetic analysis by the Maximum Likelihood (ML) method with the Le & Gascuel 2008 amino acid substitution model (Le and Gascuel 2008). The optimal tree with the highest log likelihood is shown along with the percentage of trees in which the associated taxa clustered together in the bootstrap test of 500 replicates. A discrete Gamma distribution was used to model evolutionary rate differences among sites (5 categories). Branch lengths are measured in the number of substitutions per site. All positions with less than 70% site coverage were eliminated leaving a total of 635 positions in the final dataset. (**B**). Phylogenetic analysis using the Neighbor-Joining method (Saitou and Nei 1987) gives the same result as in **A**. The percentages from 500 bootstrap replicate trees in which the associated taxa clustered together are shown next to the branches. Evolutionary distances were computed using the JTT matrix-based method (Jones *et al.* 1992) and are in the units of the number of amino acid substitutions per site. The rate variation among sites was modeled with a gamma distribution (shape parameter = 1). All positions containing gaps or missing data were eliminated leaving a total of 580 positions in the final dataset. Both the ML and NJ trees were computed using a ClustalW-based alignment of the concatenated peptide sequences from the H2B–H2A, H4–H3, and H2A-domain-containing (H2ADC) genes found in all MV genomes, and the MEGA6 software package (Tamura *et al.* 2013).

**Figure S2.**
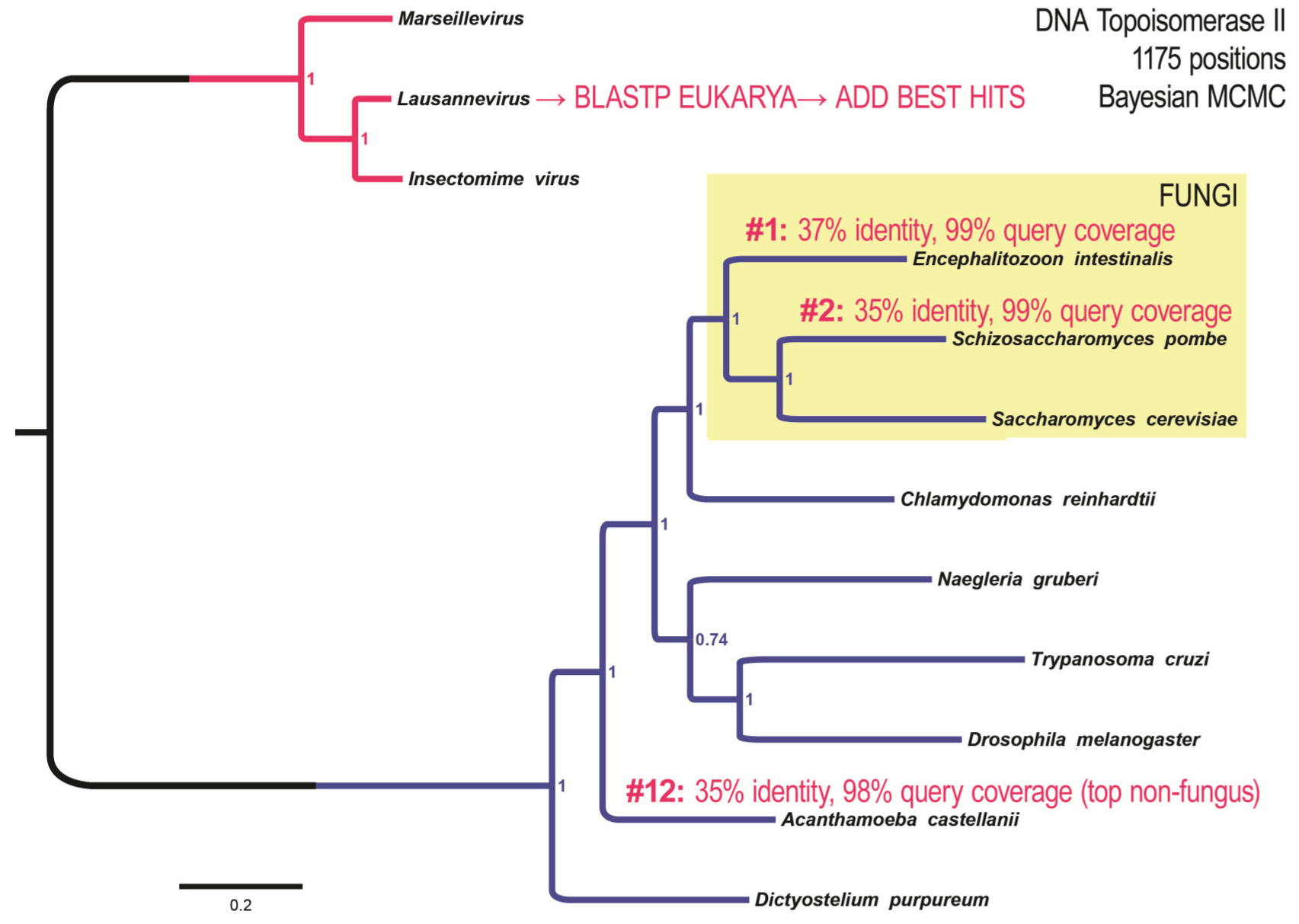
The MV DNA topoisomerase II is unassignable to any one eukaryotic lineage. The DNA topoisomerase II protein encoded by MV genomes possesses additional domains not seen in the Euarchaeota DNA gyrase subunit. To determine whether the MV DNA topoisomerase II was derived from a particular eukaryotic lineage, a Bayesian phylogenetic analysis (1M generations, 25% burn-in of sampling mixed amino acid models, with Wag having complete posterior probability and avg. st. dev. of split freq. = 0.00047) was conducted using top eukaryotic hits from a query using the Lausannevirus DNA topoisomerase II sequence. This identified fungal sequences as the top hits. The top fungal hits (labeled by % identity and % query coverage from the BLASTP query) and the top non-fungal hits were included in the analysis without the archaeal sequences. The results of this analysis are consistent with a model in which the MV DNA topoisomerase II gene was derived from an unknown stem-eukaryotic (non-eukaryotic) lineage.

